# *Bifidobacterium* is enriched in gut microbiome of Kashmiri women with polycystic ovary syndrome

**DOI:** 10.1101/718510

**Authors:** Saqib Hassan, Marika A Kaakinen, Harmen Draisma, Mohd Ashraf Ganie, Aafia Rashid, Zhanna Balkhiyarova, George Seghal Kiran, Paris Vogazianos, Christos Shammas, Joseph Selvin, Athos Antoniades, Ayse Demirkan, Inga Prokopenko

## Abstract

Polycystic ovary syndrome (PCOS) is a common endocrine condition in women of reproductive age understudied in non-European populations. In India, PCOS affects the life of up to 19.4 million women of age 14-25 years. Gut microbiome composition might contribute to PCOS susceptibility. We profiled the microbiome in DNA isolated from faecal samples by 16S rRNA sequencing in 19/20 women with/without PCOS from Kashmir, India. We assigned genera to sequenced species with an average 121k reads depth and included bacteria detected in at least 1/3 of the subjects or with average relative abundance ≥0.1%. We compared the relative abundances of 40/58 operational taxonomic units in family/genus level between cases and controls, and in relation to 33 hormonal and metabolic factors, by multivariate analyses adjusted for confounders, and corrected for multiple testing. Seven genera were significantly enriched in PCOS cases: *Sarcina, Alkalibacterium* and *Megasphaera*, and previously reported for PCOS *Bifidobacterium, Collinsella, Paraprevotella* and *Lactobacillus*. We identified significantly increased relative abundance of *Bifidobacteriaceae* (median 6.07% vs. 2.77%) and *Aerococcaceae* (0.03% vs. 0.004%), whereas we detected lower relative abundance *Peptococcaceae* (0.16% vs. 0.25%) in PCOS cases. For the first time, we identified a significant direct association between butyrate producing *Eubacterium* and follicle-stimulating hormone levels. We observed increased relative abundance of *Collinsella* and *Paraprevotella* with higher fasting blood glucose levels, and *Paraprevotella* and *Alkalibacterium* with larger hip and waist circumference, and weight. We show a relationship between gut microbiome composition and PCOS linking it to specific reproductive health metabolic and hormonal predictors in Indian women.

## Introduction

Polycystic ovary syndrome (PCOS) is a common endocrine condition affecting women of reproductive age, characterized by hyperandrogenism, oligo- or amenorrhea, and polycystic ovaries on transabdominal ultrasonography. The worldwide prevalence of PCOS among women of fertile age is about 6-18%, and it varies with the use of different diagnostic criteria of PCOS such as the Rotterdam criteria, 2003, or National Institutes of Health (NIH), the European Society of Human Reproductive and Embryology (ESHRE) and the Androgen Excess Society (AES) criteria, 2006 [1-3]. In India, PCOS affects the life of an estimated 6.5 to 19.4 million women of age 14 to 25, and the prevalence is increasing in parallel with the obesity epidemic [4]. In the 1930-s, PCOS was defined as a gynaecological disorder [5], however, it is associated with a constellation of metabolic conditions, such as obesity, dyslipidaemia, metabolic syndrome, endothelial dysfunction, inflammation, insulin resistance, hypertension and other cardiovascular risks [6-12]. Additionally, infertility, pregnancy complications and depression are frequent complications in patients with PCOS [13-15]. While the development of PCOS is multifaceted and involves genetic [16, 17], gestational environment [18] and lifestyle aspects [19]. The precise factors responsible for these key biochemical and metabolic derangements and affecting our surrounding environment as well as internal ecosystems, represented by microbiota among others, remain largely unexplored.

From genetic point of view, the human gut microbiome is a collection of microbial genomes of microorganisms that inhabit the human gut [20]. The human gut microbiome is a complex ecosystem harbouring numerous microbes taking part in essential functions of the host organism [21]. Earlier studies in European and Chinese case and control designs showed differences in diversity and relative abundance of particular taxa. It was shown that relative abundances of bacteria belonging to *Bacteroides, Escherichia/Shigella* and *Streptococcu*s, were inversely correlated with ghrelin, and positively correlated with testosterone and BMI in Chinese women with PCOS. In addition, relative abundances of *Akkermansia* and *Ruminococcaceae* are inversely correlated with body-weight, sex-hormone, and brain–gut peptides, and are decreased in PCOS [22]. The gut microbiota alterations in humans are reportedly associated with obesity [23, 24]. Additionally, the peripheral insulin sensitivity in metabolic syndrome subjects improved upon transfer of stool from healthy donors, thus suggesting a relationship between the glucose metabolism and gut microbiome [25]. Importantly, gut microbiota and its metabolites can control inflammatory processes, brain gut peptide secretion as well as islet b-cell proliferation hence may lead to excessive accumulation of fat ultimately causing insulin resistance and compensatory hyperinsulinemia [25, 26]. In a female mouse model, the testosterone levels increased upon infecting it with male faecal microbiota compared to unmanipulated females [27]. Host’s estrus cycles, sex hormones and morphological changes in ovaries are affected by gut microbial composition of the host [28]. In rats, the pre-natal exposure to high androgen levels in daughters from mothers with PCOS led to dysbiosis in gut microbiome and impairment of the cardiometabolic functions [29]. The association between gut microbiome, obesity and host genetics suggests that certain bacteria predisposing to a healthy or unhealthy metabolic state may be heritable [30-32]. It is hypothesized that diet might induce bacterial dysbiosis thus leading to inflammation, insulin resistance and hyperandrogenemia in PCOS [33].

In this study, we profiled the gut microbiome of women with PCOS and healthy controls from Northern India. We dissected the gut microbiome composition in relation to blood biochemistry and hormonal levels. Our study suggests that there are differences in the microbial composition between women with PCOS and healthy controls; two families and seven genera are significantly enriched in PCOS, whereas one family is underrepresented in women with PCOS.

## Material and Methods

### Study sample

Twenty women with PCOS (drug naive) and 20 control women without PCOS, both groups in age ranging 16-25 years, were recruited at the Endocrine clinic of the Sher-i-Kashmir Institute of Medical Sciences (SKIMS, a tertiary care hospital), Kashmir, North India from January to May 2017. The case group consisted of participants with menstrual disturbances including oligomenorrhea (menstrual interval > 35 days or < 8 cycles/year) or amenorrhea (no menstrual cycle in last >6 months), hyperandrogenism (male pattern hair growth, androgenic alopecia), polycystic ovaries on transabdominal ultrasonography and qualified Rotterdam 2003 Criteria [1] for PCOS diagnosis. The healthy controls (non-PCOS) were women having regular cyclicity in their menstrual cycles (21-35 days), no signs of hyperandrogenism and had normal ovarian morphology as evidenced through transabdominal ultrasonography. Women on antibiotic treatment or were taking contraceptives, steroids, anti-epileptics, insulin sensitizers, proton-pump inhibitors, or had any previous history of systemic sickness such as diabetes mellitus, coronary artery disease, non-classical congenital adrenal hyperplasia (NCAH), Cushing syndrome, hyperprolactinemia, thyroid dysfunction, gastrointestinal disease and appendectomy were excluded from the study. The study protocol was approved by the Institutional Ethics Committee (IEC) of SKIMS Kashmir and written informed consent was obtained from all the participants involved in this study.

### Blood biochemistry and hormonal levels

Fasting blood samples were collected in dry and EDTA coated tubes after 10-12 h fasting. Sera were separated to be used in measuring glucose, lipids, alkaline phosphatase (ALP), aspartate aminotransferase (AST), albumin, creatinine, in addition to hormones (prolactin, thyroid-stimulating hormone (TSH), thyroxine (T4), triiodothyronine (T3), cortisol, luteinizing hormone (LH), follicle-stimulating hormone (FSH), 17α-hydroxyprogesterone (17α-OHP), and total testosterone). The samples for LH, FSH, TESTO, and 17α-OHP were collected on days 3rd-7th (early follicular phase) of spontaneous cycle. The EDTA-containing aliquot was immediately placed on ice and centrifuged within 30 min; plasma was collected and was stored at −80°C until analysis for further analysis. ALP, AST, creatinine, albumin and lipid measurements were performed using fully automated chemistry analyser (Hitachi 920 at SKIMS). Estimation of serum T3, T4, cortisol, 17α-OHP, TSH, prolactin, LH, FSH and total T4 was done by RIA using commercial kits and according to supplier protocol at SKIMS. Plasma glucose was measured by glucose oxidase peroxidase method.

### Stool samples

Each PCOS patient and healthy participant (non-PCOS) was asked to provide fresh stool sample (approx. 5 grams) within the same week when the blood was collected by using the stool collection and stabilization kit (OMNIgene®•GUT OMR-200, DNA Genotek, Canada) which was given to all the participants. All samples were collected in the morning and were stored at ambient temperature until further processing which took place within the next 60 days. Consistency of each sample was recorded on the following scale: 1= Sausage shaped with cracks on the surface, 2 =Sausage shaped and smooth soft stool, 3= Solid stool with clumps, 4= Watery stool

### Metagenome DNA extraction and 16S rRNA sequencing data generation

DNA extraction was performed by as per the instruction manual using ZymoBIOMICS™ DNA kit by Zymo Research USA. The DNA concentrations were estimated by Qubit Fluorometer (Thermo Fisher) and checked by Agilent TapeStation 2200. The microbiota characterization was performed by targeting the hypervariable regions V3-V4 of 16S rRNA gene using paired-end approach using the specific primers published earlier [34] and according to the manufacturer’s instructions. [35] The amplified regions were combined with dual-index barcodes, Nextera® XT Index Kit v2 Set A, B and C, Illumina USA. The sequencing run was performed with MiSeq 600 cycle Reagent Kit v3, Illumina USA and sequenced on a MiSeq Illumina system by the Stremble Ventures LTD, Cyprus.

### Bioinformatics and quality control

The bioinformatic analysis was performed from raw FASTQ files with paired-end reads created classified operation taxonomic units (OTU) using Ribosomal Database Project Classifier (version/access date) [36] against the Illumina-curated version of GreenGenes reference taxonomy database (version/access date) [37]. OTUs that were detected in more than 1/3 of the sample were considered prevalent OTUs and among those, the ones with > 0.1 % average relative abundance across all subjects were included in analysis.

### Statistical analysis

#### Microbial distance and diversity

Inter-individual microbial distance (Bray-Curtis distance) and diversity (Shannon’s diversity index) were calculated using functions *vegdist* and *diversity* the R package *vegan* [38]. We analysed the association between PCOS and i) Bray-Curtis distance using the function *adonis* in R package *vegan*, and ii) Shannon’s diversity index using Spearman correlation in R. Both analyses were additionally adjusted for stool consistency, day of menstrual cycle at sample collection and sequencing read depth.

#### Association analyses between individual species, PCOS and hormone levels

We performed all the analyses at Genus and Family levels for which the proportion of successfully classified reads across samples was over 95%. We used the non-parametric Mann-Whitney U-test [39] to compare median abundancies of each species between PCOS cases and controls. We considered results with a *P*-value<0.05 as statistically significant.

To account for potential confounding factors, we used the Multivariate Association with Linear Model, MaAsLin, in R [40], which allows for the use of covariates in the model. In brief, MaAsLin fits a linear model for each species and the variable of interest, here PCOS, after arcsin-square-root transformation of the proportional values of the species. This transformation has been shown to stabilise variance and normalise proportional data well [40]. The method can also include a boosting step to select factors among a large set of variables to be associated with the species. We turned this feature off since we focused only on one variable of interest, PCOS. We used the default settings of the programme, i.e. minimum relative abundance of 0.01%; minimum percentage of samples in which a feature must have the minimum relative abundance in order not to be removed of 10%; outlier removal by Grubbs test with the significance cut-off used to indicate an outlier at 0.05; multiple test correction by the Benjamini-Hochberg (BH) [41]method; and the threshold to use for significance for the generated q-values (BH FDR) of 0.25. As potential confounders in the model we used stool consistency, read depth, age, day of the menstrual cycle during sample collection and body mass index (BMI).

The linear model analysis with MaAsLin was performed also for LH, FSH, and testosterone levels in cases and controls together, adjusting for stool consistency, read depth, age, day of the menstrual cycle during sample collection and BMI.

For bacteria that showed differential enrichment we performed follow-up analyses to dissect the mechanistic links with PCOS. For this, we used MaAsLin to perform linear regression of the relative abundancies in relation to relevant variables that were available (**Supplementary Table 1**), including blood glucose levels and BMI. We used the same adjustments as before.

**Table 1.** Characteristics of the study sample.

|  | Cases (N=19) | Controls (N=20) | P-value* |
| --- | --- | --- | --- |
|  | Mean (SD) or<br>N[%] | Mean (SD) or N[%] |  |
| **Anthropometrics and blood measurements** |  |  |  |
| Age, years | 23.9 (6.9) | 21.1 (2.5) | 0.093 |
| Height, cm | **155.6 (4.9)** | **151.5 (6.8)** | **0.037** |
| Weight, kg | **61.5 (8.9)** | **53.1 (7.5)** | **2.86E-03** |
| BMI, kg/m<sup>2</sup> | **25.4 (3.3)** | **23.2 (3.2)** | **0.041** |
| Waist, cm | **81.3 (10)** | **66.4 (4.7)** | **6.00E-07** |
| Hip circumference, cm | **92.2 (7.8)** | **75.9 (5.3)** | **3.63E-09** |
| Waist-hip-ratio | 0.9 (0.1) | 0.9 (0) | 0.73 |
| Systolic blood pressure, mmHg | 124.5 (9.7) | 120.8 (6.1) | 0.16 |
| Diastolic blood pressure, mmHg | 81.6 (7.6) | 80.5 (3.9) | 0.58 |
| Luteinizing hormone (LH), IU/L | 5 (2.8) | 4.3 (0.6) | 0.28 |
| Follicle stimulating hormone (FSH),<br>IU/L | 5.5 (1.6) | 5.1 (1.2) | 0.34 |
| LH/FSH ratio | 0.97 (0.70) | 0.90 (0.28) | 0.66 |
| Testosterone, ng/dL | 56.8 (26.6) | 49.6 (12.7) | 0.28 |
| Prolactin | **15.7 (7.2)** | **10 (4.7)** | **5.64E-03** |
| Fasting blood glucose, mg/dL | **114.2 (8.6)** | **106.1 (9.1)** | **6.71E-03** |
| Blood glucose 1 hour, mg/dL | 126 (11.8) | 120 (24.3) | 0.33 |
| Blood glucose 2 hour, mg/dL | 121.2 (10.7) | 121.3 (22.3) | 0.99 |
| Total cholesterol, mg/dL | 164.7 (31.8) | 149.1 (26.9) | 0.11 |
| Triglycerides, mg/dL | 205.9 (105.8) | 144.3 (96.6) | 0.065 |
| High-density lipoprotein (HDL), mg/dL | 39.6 (4.6) | 41.8 (4.4) | 0.14 |
| Low-density lipoprotein (LDL), mg/dL | 110.2 (8.4) | 108.4 (9.5) | 0.54 |
| **PCOS-related** |  |  |  |
| Age at menarche, years | 13.3 (2.2) | 12.2 (1.2) | 0.063 |
| Number of cycles per year | 6.8 (1.9) | 10.7 (1) | 1.69E-09 |
| H-score | 7.3 (5.5) | 0.4 (1.8) | 6.03E-06 |
| Menstrual cycle irregularity, yes | 19 [100] | 0 [0] | 1.45E-11 |
| Acanthosis nigricans, yes | 5 [26] | 0 [0] | 0.02 |
| Acne, yes | 7 [37] | 2 [10] | 0.06 |
| Alopecia, yes | 9 [47] | 4 [20] | 0.10 |
| Duration of hirsutism, years | 1.8 (0.8) | 0.1 (0.2) | 1.87E-11 |
| Family history of hirsutism, yes | 3 [16] | 2 [10] | 0.66 |
| Family history of menstrual disturbances, yes | 5 [26] | 2 [10] | 0.24 |
| Family history of T2D, yes | 11 [58] | 7 [35] | 0.2 |
| **Stool sample collection and sequencing** |  |  |  |
| Day of menstrual cycle at stool collection |  |  |  |
| Day 3 | 7 [37] | 6 [30] | 1.00 |
| Day 4 | 5 [26] | 6 [30] |  |
| Day 5 | 6 [32] | 7 [35] |  |
| Day 6 | 1 [5] | 1 [5] |  |
| Stool consistency |  |  |  |
| Sausage shaped with cracks on the surface | 3 [16] | 2 [10] | 0.66 |
| Sausage shaped and smooth soft stool | 12 [63] | 15 [75] |  |
| Solid clumpy stool | 3 [16] | 1 [5] |  |
| Watery stool | 1 [5] | 2 [10] |  |
| Read depth, genus level | 80434.6 (44927.2) | 161279.9 (158861.1) | 0.039 |
| Read depth, family level | 81547.8 (45539.7) | 164695 (162499.2) | 0.038 |

\*P-value is from t-test for continuous traits and Fisher's exact test for categorical traits.

## Results

### 16S rRNA gut microbiome in PCOS

After the quality control of 16S rRNA sequencing, we included in the analysis genetic data for 39 microbiomes of Kashmiri women with/without PCOS (**Table 1, Supplementary Table 1**), including 58 OTUs in the genus level and 40 in the family level, detected in at least 30% of the samples with an average frequency of 0.1%.

### Associations with PCOS at genus and family level

The analyses of the individual species at genus level showed statistically significant (*P*<0.05) differences between cases and controls for 12 organisms (**Table 2**). The most striking differences were observed for *Sarcina* (cases vs controls: 0.28% vs. 0.06%), *Megasphaera* (3.62% vs. 1.17%) and *Bifidobacterium* (7.7% vs. 3.1%) (**Figure 1**). Nine of these bacteria reached FDR corrected statistical significance also in the linear modelling with arcsin-square root transformation and outlier exclusions, with *Sarcina, Alkalibacterium, Megasphaera, Collinsella, Paraprevotella, Lactobacillus* and *Bifidobacterium* surviving adjustment for stool consistency, day of menstrual cycle at data collection point, sequencing read depth and age (**Table 2, Supplementary Table 7**). When the model was additionally adjusted for BMI, *Lactobacillus* did not reach the FDR-corrected level of significance anymore.

**Table 2.** Statistically significant differences between PCOS cases and controls in the relative abundancies at genus level from the Mann-Whitney U-test (*P*<0.05) or from the multivariate model (Q<0.25). Values highlighted in bold show significant associations in either test and within each adjustment.

|  | Mann-Whitney U-test |  |  | MaAsLin |  |  |  |  |  |  |  |  |  |
| --- | --- | --- | --- | --- | --- | --- | --- | --- | --- | --- | --- | --- | --- |
|  |  |  |  | Unadjusted |  |  |  | Adjusted for stool consistency, read depth, day of menstrual cycle at sample collection, age |  |  | Adjusted for stool consistency, read depth, day of menstrual cycle at sample collection, age, BMI |  |  |
| Feature | Mean (%) in cases (N=19) | Mean (%) in controls (N=20) | P-value | N | Coefficient | P-value | Q-value | Coefficient | P-value | Q-value | Coefficient | P-value | Q-value |
| *Sarcina* | **0.28** | **0.059** | **$3.4 \times 10^{-4}$** | 36 | **0.023** | **0.001** | **0.025** | **0.023** | **0.001** | **0.066** | **0.024** | **0.001** | **0.127** |
| *Megasphaera* | **3.62** | **1.17** | **$2.1 \times 10^{-3}$** | 39 | **0.093** | **0.003** | **0.048** | **0.099** | **0.003** | **0.104** | **0.098** | **0.005** | **0.190** |
| *Bifidobacterium* | **7.72** | **3.12** | **$7.5 \times 10^{-3}$** | 39 | **0.096** | **0.006** | **0.048** | **0.100** | **0.004** | **0.115** | **0.103** | **0.005** | **0.190** |
| *Paraprevotella* | **0.18** | **0.032** | **$7.3 \times 10^{-3}$** | 37 | **0.014** | **0.004** | **0.048** | **0.014** | **0.001** | **0.066** | **0.012** | **0.006** | **0.192** |
| *Collinsella* | **1.68** | **0.78** | **$7.5 \times 10^{-3}$** | 39 | **0.041** | **0.004** | **0.048** | **0.043** | **0.002** | **0.101** | **0.043** | **0.004** | **0.179** |
| *Erysipelothrix* | **1.52** | **0.29** | **$7.5 \times 10^{-3}$** | 39 | **0.017** | **0.036** | **0.219** | 0.018 | 0.022 | 0.342 | 0.016 | 0.058 | 0.475 |
| *Lactobacillus* | **5.30** | **1.90** | **0.012** | 39 | **0.086** | **0.006** | **0.048** | **0.085** | **0.008** | **0.207** | 0.084 | 0.014 | 0.304 |
| *Dysgonomonas* | **0.66** | **0.18** | **0.013** | 39 | **0.016** | **0.035** | **0.219** | 0.016 | 0.047 | 0.481 | 0.016 | 0.063 | 0.501 |
| *Oscillospira* | **2.22** | **1.68** | **0.018** | 39 | 0.033 | 0.168 | 0.454 | 0.039 | 0.06 | 0.487 | 0.036 | 0.104 | 0.574 |
| *Natronincola* | **0.35** | **0.31** | **0.030** | 38 | 0.016 | 0.049 | 0.256 | 0.017 | 0.016 | 0.298 | 0.016 | 0.031 | 0.456 |
| *Alkalibacterium* | **0.089** | **0.32** | **0.049** | 35 | **0.007** | **0.001** | **0.025** | **0.007** | **0.000** | **0.066** | **0.006** | **0.002** | **0.145** |
| *Atopobium* | 0.27 | 0.08 | 0.066 | 36 | **0.020** | **0.038** | **0.219** | 0.019 | 0.042 | 0.481 | 0.019 | 0.057 | 0.475 |

**Figure 1.**
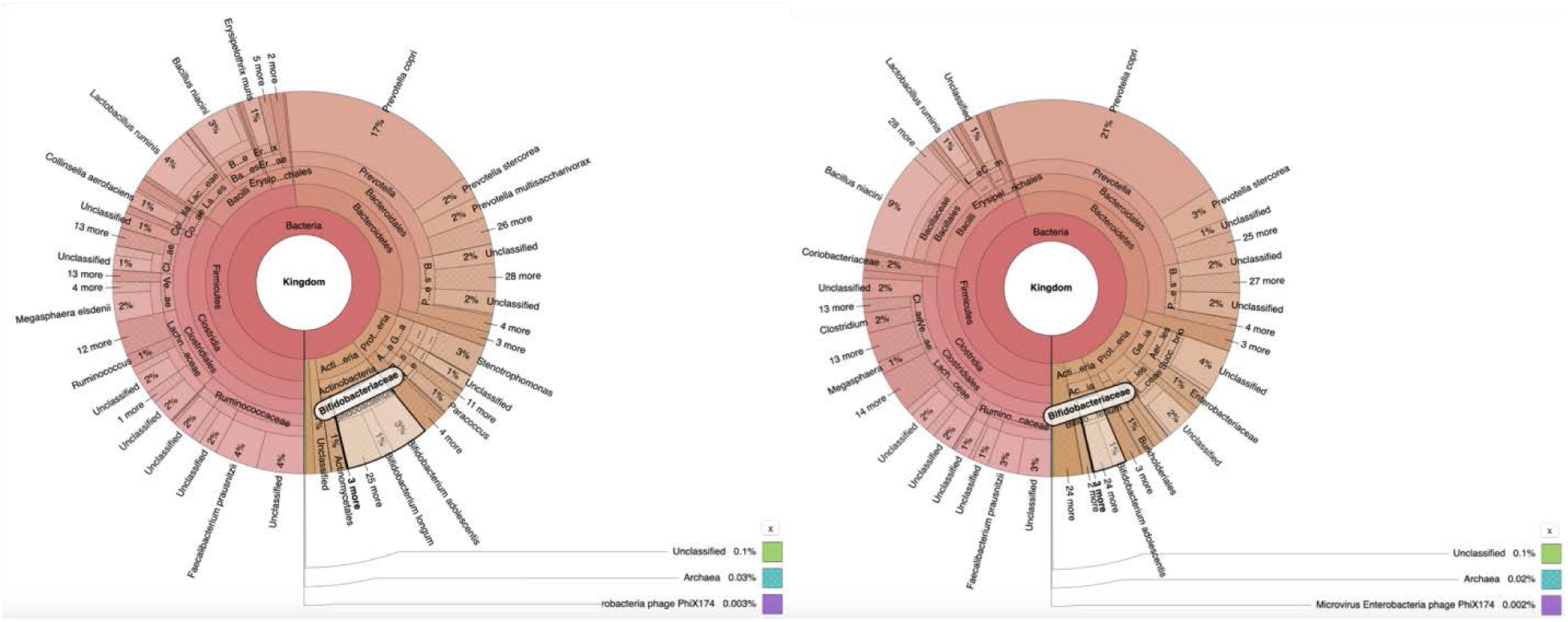
Krona plots showing the bacterial composition in PCOS A) cases, and B) controls. The family *Bifidobactriaceae* is highlighted as an example showing enrichment in cases as compared to controls.

Five families showed nominally significant differences between PCOS cases and controls by the Mann-Whitney U-test. These included *Peptococcaceae, Bifidobacteriaceae, Lactobacillaceae, Erysipelotrichaceae* and *Porphyromonadaceae* (**Table 3**). The linear modelling with MaAslin confirmed the associations with *Peptococcaceae* and *Bifidobacteriaceae*, and showed additionally a significant association with *Aerococcaceae* (**Figures 2** and **3, Supplementary Figure 1**). The association with *Lactobacillaceae* did not survive the adjustments for stool consistency, day of menstrual cycle at data collection point, sequencing read depth and age (**Table 3, Supplementary Table 2**).

**Table 3.** Statistically significant differences between PCOS cases and controls in the relative abundancies at family level from the Mann-Whitney U-test (*P*<0.05) or from the multivariate model (Q<0.25). Values highlighted in bold show significant associations in either test and within each adjustment.

| Feature | Mann-Whitney U-test |  |  | MaAsLin |  |  |  |  |  |  |  |  |  |
| --- | --- | --- | --- | --- | --- | --- | --- | --- | --- | --- | --- | --- | --- |
|  | Mean (%) | Mean (%) | P-value | Unadjusted |  |  |  | Adjusted for stool consistency, read depth, day of menstrual cycle at sample collection, age |  |  | Adjusted for stool consistency, read depth, day of menstrual cycle at sample collection, age, BMI |  |  |
|  | in cases (N=19) | in controls (N=20) |  |  |  |  |  |  |  |  |  |  |  |
|  |  |  |  | N | Coefficient | P-value | Q-value | Coefficient | P-value | Q-value | Coefficient | P-value | Q-value |
| *Peptococcaceae* | **0.15** | **0.27** | **$1.5 \times 10^{-3}$** | 39 | **-0.011** | **0.003** | **0.055** | **-0.010** | **0.005** | **0.246** | **-0.010** | **0.005** | **0.233** |
| *Bifidobacteriaceae* | **7.63** | **3.07** | **$7.5 \times 10^{-3}$** | 39 | **0.096** | **0.005** | **0.055** | **0.100** | **0.003** | **0.233** | **0.103** | **0.005** | **0.233** |
| *Lactobacillaceae* | **5.26** | **1.89** | **0.012** | 39 | 0.086 | 0.005 | 0.055 | 0.085 | 0.008 | 0.307 | 0.083 | 0.014 | 0.349 |
| *Erysipelotrichaceae* | **2.50** | **0.91** | **0.026** | 39 | 0.030 | 0.034 | 0.254 | 0.029 | 0.045 | 0.457 | 0.022 | 0.124 | 0.680 |
| *Porphyromonadaceae* | **0.88** | **0.43** | **0.047** | 39 | 0.015 | 0.058 | 0.254 | 0.016 | 0.051 | 0.461 | 0.015 | 0.094 | 0.680 |
| *Aerococcaceae* | 0.08 | 0.31 | 0.052 | **35** | **0.007** | **0.001** | **0.037** | **0.007** | **0.000** | **0.082** | **0.006** | **0.002** | **0.233** |

**Figure 2.**
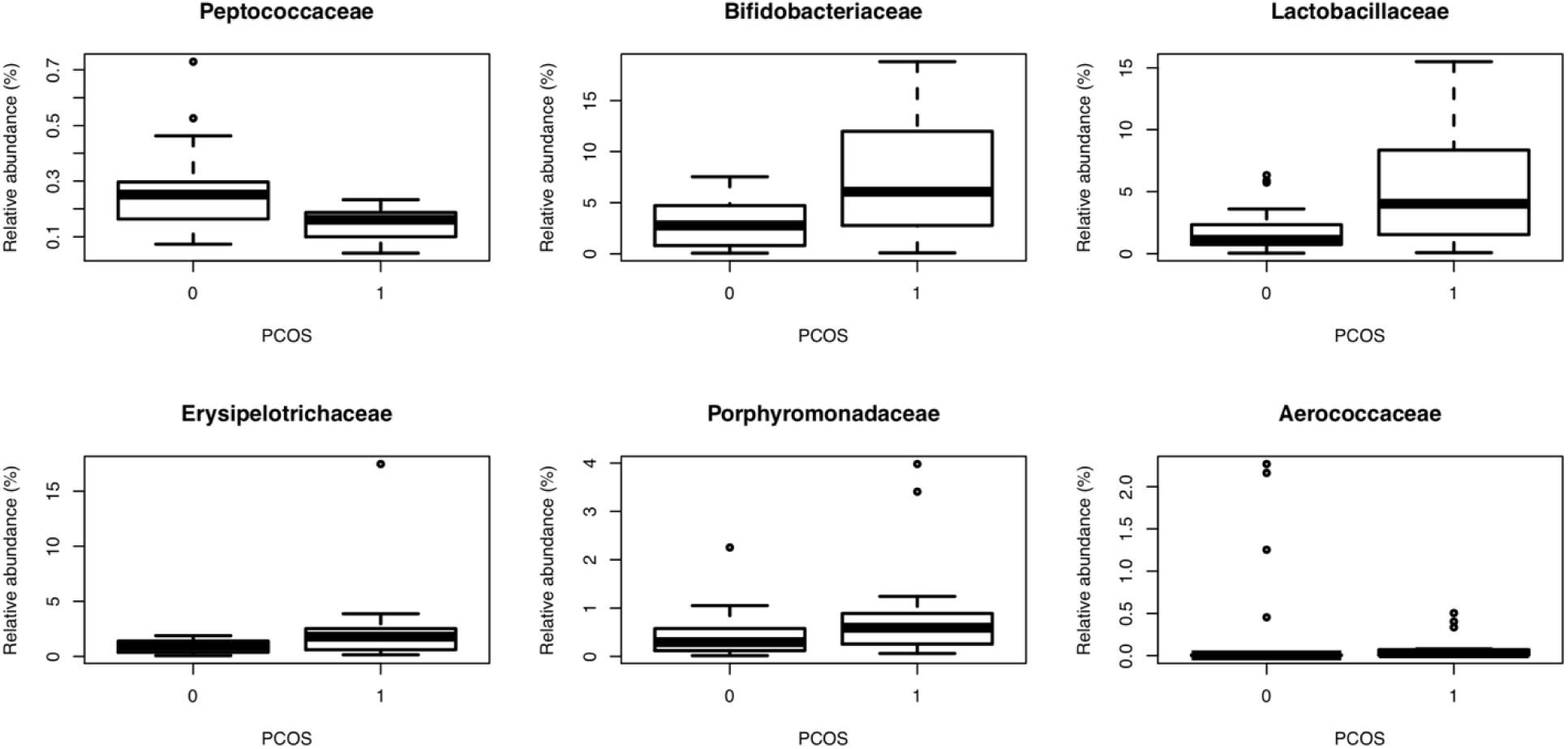
Boxplots showing the distributions of the six bacterial families reaching statistical significance either in the Mann-Whitney U-test for PCOS cases and controls (P<0.05) or in the linear modelling with MaAsLin (Q<0.25).

**Figure 3.**
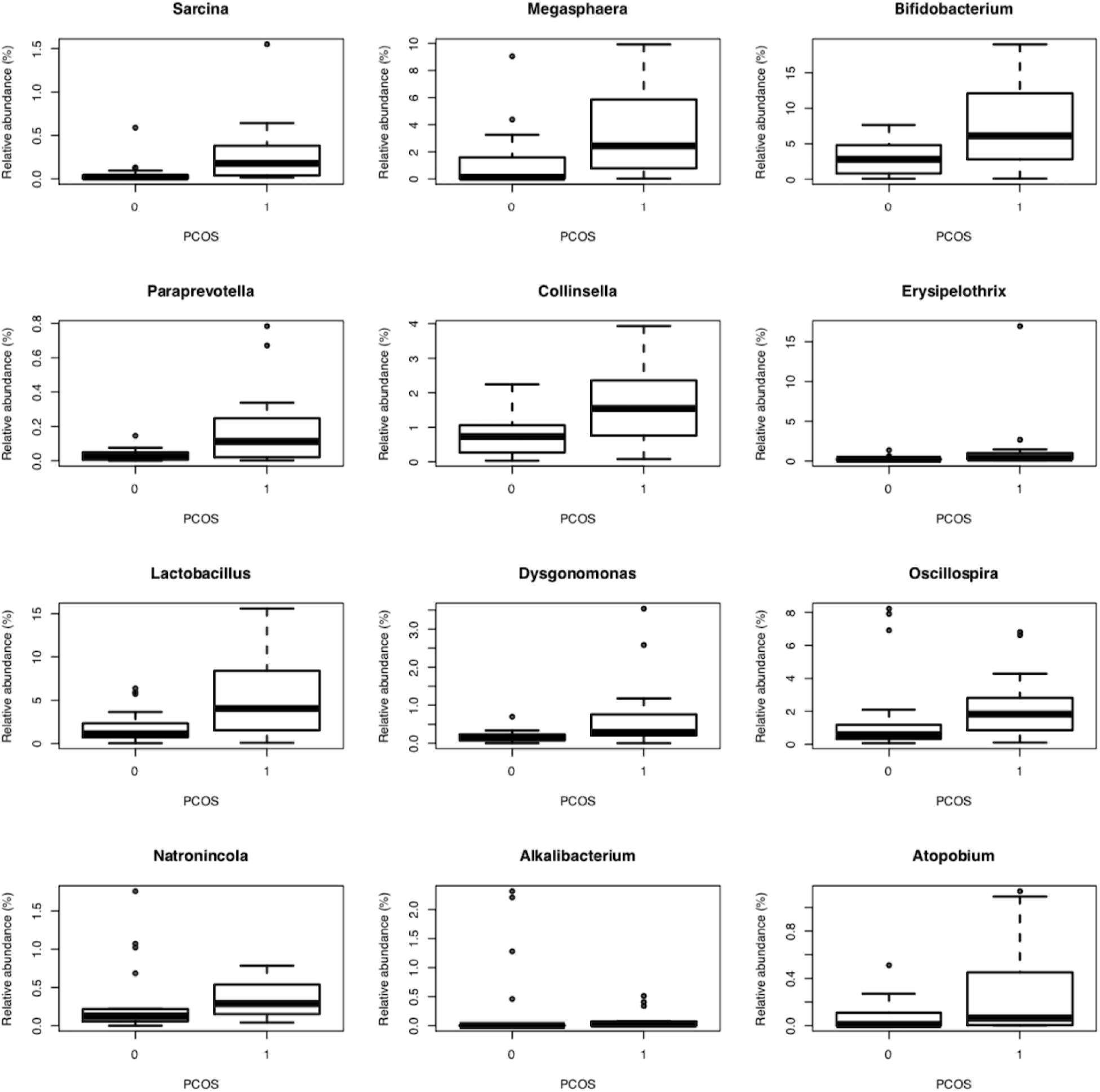
Boxplots showing the distributions of the 12 bacterial species reaching statistical significance either in the Mann-Whitney U-test for PCOS cases and controls (*P*<0.05) or in the linear modelling with MaAsLin (Q<0.25).

### Quantitative traits and hormonal profiles at genus and family level

The cases were taller, had higher weight, BMI, waist and hip circumference than the controls (**Table 1**). They also had higher blood glucose, ALP, albumin, creatinine, prolactin and TSH levels and lower AST levels (**Table 1, Supplementary Table 1**).

The quantitative trait analyses detected association between higher fasting blood glucose levels and enrichment in *Collinsella* (0.003, *P*=3.4×10^−4^, Q=0.16) and *Paraprevotella* (0.001, *P*=1.4×10^−3^, Q=0.17). Additionally, we identified enrichment for *Paraprevotella* with larger hip (0.001, *P*=2.9×10^−3^, Q=0.20), waist circumference (0.001, *P*=1.0×10^−3^, Q=0.17) and weight (0.001, *P*=2.1×10^−3^, Q=0.20). Similar enrichment was detected in *Alkalibacterium* for hip circumference (0.0003, *P*=2.5×10^−3^, Q=0.20), waist circumference (0.0003, *P*=1.4×10^−3^, Q=0.17), and weight (0.0004, *P*=1.7×10^−4^, Q=0.19).

Upon linear modelling for the hormonal profiles (FSH, LH, LH to FSH and testosterone) of the study participants, instead of the binary case control status, *Eubacterium* reached the FDR-corrected significance level (Q<0.25) for association with FSH. This association remained significant after adjustment for the covariates (**Supplementary Table 7**). Our follow-up analyses with other phenotypes indicated that *Eubacterium* was associated with weight (0.003, *P*=3.8×10^− 4^, Q=0.16). For LH, LH to FSH and testosterone, we observed no statistically significant associations for any of the bacterial abundancies (**Supplementary Tables 8-11**).

At family level, we detected an association between increased *Peptococcaceae* and lower prolactin levels (−0.001, *P*=4.2×10^−4^, Q=0.19). Linear regression analysis of hormonal levels and family level relative abundancies did not yield any statistically significant associations (**Supplementary Tables 3-6**).

### Distance and diversity

The correlation between bacterial alpha-diversity was statistically different in PCOS cases compared to the controls (r=0.40, *P*=0.012, **Figure 4A**) when adjusting for stool consistency, day of menstrual cycle at sample collection, and read depth. However, the bacterial diversity was not significantly different in the two studied groups (r=0.22, *P*=0.18) without using the above adjustments (**Figure 4B**). Similarly, we observed no statistically significant differences neither in the unadjusted nor adjusted bacterial compositions between cases and controls as measured by the Bray-Curtis distance (unadjusted *P*=0.10, adjusted *P*=0.09). Both at genus and family level, the sequencing yielded a higher mean number of reads in controls as compared to the cases (**Table 1**).

**Figure 4.**
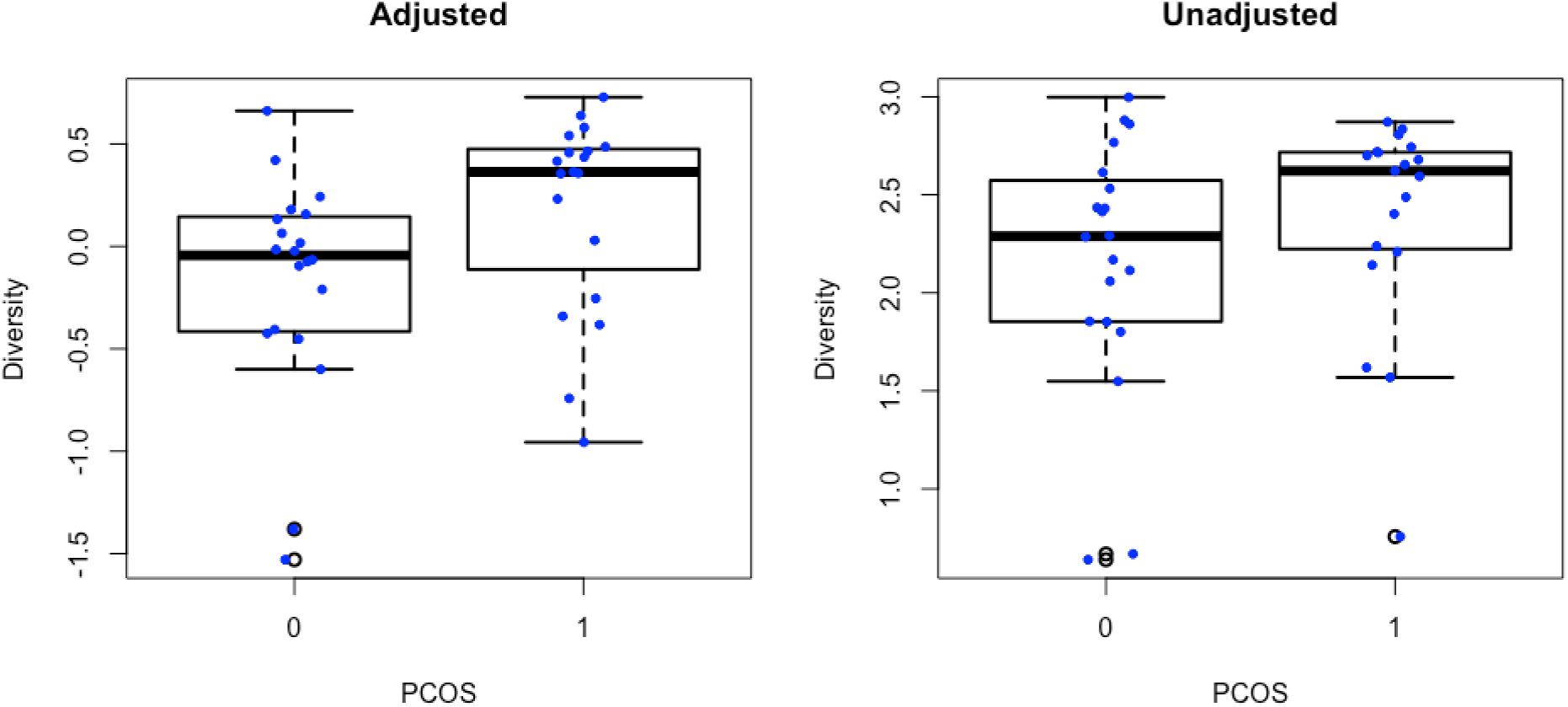
Boxplots showing the diversity in bacterial composition measured as Shannon’s diversity index between PCOS cases and controls. A) Adjusted for stool consistency, day of menstrual cycle at sample collection and sequencing read depth, B) Unadjusted.

## Discussion

This is a first investigation of PCOS gut microbiome in women from Kashmir, India. Here, we compared 39 individuals with/without PCOS by their gut microbiome composition and dissected the latter in relation to 33 quantitative endophenotypes. We performed a step-wise association modelling moving from simplistic towards more complex covariate models and identified robust signals both with genus and family level taxonomy. Seven genera and three family-level groups were significantly enriched in PCOS cases, including enrichment at both levels in *Bifidobacterium and Bifidobacteriaceae*. For the first time, we identified a significant direct association between butyrate producing *Eubacterium* and follicle-stimulating hormone levels.

Overall, the gut microbiome of women with PCOS has a higher bacterial diversity compared to that of women without PCOS, when measured by Shannon’s index. However, women with PCOS also have lower bacterial gene count. In addition, for multiple OTUs, the PCOS cases show higher relative abundancies when compared to the healthy controls.

At the genus level, relative abundances of seven genera including *Lactobacillus, Bifidobacterium, Sarcina, Alkalibacterium, Megasphaera, Collinsella* and *Paraprevotella*, are increased in gut microbiome of women with PCOS. Association of *Lactobacilius* seems to be dependent on BMI, whereas the other genera remained significantly associated after adjustment with BMI. Among those we show that *Collinsella* and *Paraprevotella* were associated with higher fasting blood glucose levels, and *Paraprevotella* and *Alkalibacterium* further associate with larger hip and waist circumference and weight. Lastly, *Eubacterium* is positively linked with FSH level. Our study is the first to report *Sarcina, Alkalibacterium* and *Megasphaera* in relation to PCOS. At the family level, the gut microbiome of women with PCOS are enriched with *Aerococcaceae* and *Bifidobacteriaceae* whereas they harbour lower relative abundance of the family *Peptococcaceae* when we account for the technical covariates in the statistical models. The associations are not likely to be confounded by the BMI of the participants.

To date, a few observational studies on the gut microbiome of individuals with PCOS exist [20, 22, 28, 42, 43]. *Bifidobacterium* intake resulted in lower insulin concentration, insulin resistance and higher insulin sensitivity in patients after a 12-week intervention[44]. Relative *Bifidobacterium* abundance in human gut is known to be driven by lactose intolerance. In lactose-intolerant individuals, lactose is not metabolized in the small intestine and proceeds to the colon where it is fermented by members of the gut microbiome; this fermentation leads to gas production, a major symptom associated with lactose intolerance [45]. Thus, genetic variants that reduce lactase activity can promote the growth of lactose-fermenting bacteria in the colon, but only if the individual consumes dairy products. Taken together with our findings this brings in mind if there is an association between adult type hypolactasia and PCOS. There has been only one small study looking at the association, however it was not replicated [46]. Bifidobacterium abundance in gut is known to associate with favorable metabolic outcomes but, a recent report also showed that not all strains of Bifidobacterium are functional [47] and these strain level differences can only be resolved by metagenomics sequencing.

*Collinsella* and *Paraprevotella* were previously reported to be increased in obese controls, in comparison to women with PCOS and non-obese controls in a study with limited sample size indicating that these genera could be more specific to obesity and insulin resistance than the PCOS phenotype [22]. In contradiction with our finding, *Lactobacillius*, for which we found the association to depend on BMI, was found to be enriched in non-obese controls in comparison to the obese PCOS group in the same study. A randomized controlled study focusing of metabolic benefits synbiotics in PCOS reported that *Lactobacillus* intake resulted in lower insulin concentration, insulin resistance and higher insulin sensitivity[44] The genera *Alkalibacterium* and *Megasphaera* which we found enriched in PCOS samples are also involved in lactic acid fermentation otherwise unknown significance in human gut. *Sarcina* is a member of the family *Clostridiaceae* and species from this genus *Sarcina* ventriculi is an increasingly common gram-positive coccus, recognized in gastric biopsies, particularly of patients with delayed gastric emptying [48].

*Eubacterium* genus, associated with FSH in our study, contains a prominent human gut symbiont species with its own specific and competitive starch utilization pathway (*Eubacterium rectale*), but also contains butyrate producing bacteria such as *E*. *limosum*, in addition to several other species that fall under *Eubacterium*. Butyrate, in particular, is involved in a number of beneficial processes to the host, including downregulation of bacterial virulence; maintenance of colonic homeostasis, including acting as an energy source for intestinal epithelial cells; and anti-inflammatory effects [49].

One limitation of our study is the sample size which is comparable to the size of other pilot studies in the field. In order to deal with limited sample size, we also used the non-parametric Mann-Whitney U-test. Although this method does not allow to account for covariates, the associations with families *Peptococcaceae, Bifidobacteriaceae* and *Lactobacillaceae* and associations with genera *Sarcina, Megasphaera, Bifidobacterium, Paraprevotella* and *Collinsella* turn out to be robust, confirmed by both statistical methods. Another limitation of the study is the lack of resolution; however, this can be avoided by using shotgun metagenomics sequencing in further research. Phylogenetic profile in saliva microbiome is also altered and have lower diversity in PCOS and it is associated with clinical parameters [50]. However, none of these pilot studies were replicated at the genus level. Although association with specific taxa are not replicated yet, the emerging field of faecal microbiota transplantation shows encouraging results in rats, decreasing androgen biosynthesis and normalizing ovarian morphology [28].

We show an increased abundance of genera involved in lactic acid fermentation in the stool samples of PCOS group. Taken together with known beneficial effects of some of bacterial genera, it is difficult to conclude if these are a cause or consequence of the metabolic disturbances seen in PCOS or if they are specific to Indian women. These findings strengthen the role of gut microbiota in PCOS hormonal levels maintenance and warrants further better powered research.

## Acknowledgements

The authors would like to thank British Council, United Kingdom and Department of Biotechnology (DBT), Govt. of India for the Newton Bhabha Fellowship (BT/IN/UK/DBT-BC/2015-16). JS and GSK are thankful to Department of Biotechnology (BT/PR15632/AAQ/3/813/2016) and EU H2020 project on MicrobiomeSupport (EU: No.818116).

## References

1. Rotterdam, E.A.-S.P.c.w.g., Revised 2003 consensus on diagnostic criteria and long-term health risks related to polycystic ovary syndrome (PCOS). Hum Reprod, 2004. 19(1): p. 41–7.

2. March, W.A., et al., The prevalence of polycystic ovary syndrome in a community sample assessed under contrasting diagnostic criteria. Hum Reprod, 2010. 25(2): p. 544–51.

3. Azziz, R., et al., Positions statement: criteria for defining polycystic ovary syndrome as a predominantly hyperandrogenic syndrome: an Androgen Excess Society guideline. J Clin Endocrinol Metab, 2006. 91(11): p. 4237–45.

4. Rashid, A., et al., Evaluation of serum anti-nuclear antibody among women with PCOS: a hospital based single center cross sectional study. Gynecol Endocrinol, 2018: p. 1–5.

5. I.F., S. and L. M.L., Amenorrhea associated with bilateral polycystic ovaries. American Journal of Obstetrics and Gynecology, 1935. 29(2): p. 181–191.

6. Teede, H., A. Deeks, and L. Moran, Polycystic ovary syndrome: a complex condition with psychological, reproductive and metabolic manifestations that impacts on health across the lifespan. BMC Med, 2010. 8: p. 41.

7. Wild, R.A., et al., Lipoprotein lipids in women with androgen excess: independent associations with increased insulin and androgen. Clin Chem, 1990. 36(2): p. 283–9.

8. Dunaif, A., et al., Profound peripheral insulin resistance, independent of obesity, in polycystic ovary syndrome. Diabetes, 1989. 38(9): p. 1165–74.

9. Kelly, C.C., et al., Low grade chronic inflammation in women with polycystic ovarian syndrome. J Clin Endocrinol Metab, 2001. 86(6): p. 2453–5.

10. Repaci, A., A. Gambineri, and R. Pasquali, The role of low-grade inflammation in the polycystic ovary syndrome. Mol Cell Endocrinol, 2011. 335(1): p. 30–41.

11. Wild, R.A., et al., Assessment of cardiovascular risk and prevention of cardiovascular disease in women with the polycystic ovary syndrome: a consensus statement by the Androgen Excess and Polycystic Ovary Syndrome (AE-PCOS) Society. J Clin Endocrinol Metab, 2010. 95(5): p. 2038–49.

12. Dumesic, D.A., et al., Scientific Statement on the Diagnostic Criteria, Epidemiology, Pathophysiology, and Molecular Genetics of Polycystic Ovary Syndrome. Endocr Rev, 2015. 36(5): p. 487–525.

13. Wehr, E., et al., The lipid accumulation product is associated with impaired glucose tolerance in PCOS women. J Clin Endocrinol Metab, 2011. 96(6): p. E986–90.

14. Lerchbaum, E., et al., Assessment of glucose metabolism in polycystic ovary syndrome: HbA1c or fasting glucose compared with the oral glucose tolerance test as a screening method. Hum Reprod, 2013. 28(9): p. 2537–44.

15. Kollmann, M., et al., Maternal and neonatal outcomes in pregnant women with PCOS: comparison of different diagnostic definitions. Hum Reprod, 2015. 30(10): p. 2396–403.

16. Vink, J.M., et al., Heritability of polycystic ovary syndrome in a Dutch twin-family study. J Clin Endocrinol Metab, 2006. 91(6): p. 2100–4.

17. Shabir, I., et al., Prevalence of metabolic syndrome in the family members of women with polycystic ovary syndrome from North India. Indian J Endocrinol Metab, 2014. 18(3): p. 364–9.

18. Abbott, D.H., et al., Insights into the development of polycystic ovary syndrome (PCOS) from studies of prenatally androgenized female rhesus monkeys. Trends Endocrinol Metab, 1998. 9(2): p. 62–7.

19. Zhang, J., et al., High Intake of Energy and Fat in Southwest Chinese Women with PCOS: A Population-Based Case-Control Study. PLoS One, 2015. 10(5): p. e0127094.

20. Lindheim, L., et al., Alterations in Gut Microbiome Composition and Barrier Function Are Associated with Reproductive and Metabolic Defects in Women with Polycystic Ovary Syndrome (PCOS): A Pilot Study. PLoS One, 2017. 12(1): p. e0168390.

21. Heintz-Buschart, A. and P. Wilmes, Human Gut Microbiome: Function Matters. Trends Microbiol, 2018. 26(7): p. 563–574.

22. Liu, R., et al., Dysbiosis of Gut Microbiota Associated with Clinical Parameters in Polycystic Ovary Syndrome. Front Microbiol, 2017. 8: p. 324.

23. Turnbaugh, P.J., et al., An obesity-associated gut microbiome with increased capacity for energy harvest. Nature, 2006. 444(7122): p. 1027–31.

24. Turnbaugh, P.J., et al., A core gut microbiome in obese and lean twins. Nature, 2009. 457(7228): p. 480–4.

25. Vrieze, A., et al., Transfer of intestinal microbiota from lean donors increases insulin sensitivity in individuals with metabolic syndrome. Gastroenterology, 2012. 143(4): p. 913–6 e7.

26. Barber, T.M., et al., Polycystic ovary syndrome: insight into pathogenesis and a common association with insulin resistance. Clin Med (Lond), 2016. 16(3): p. 262–6.

27. Markle, J.G., et al., Sex differences in the gut microbiome drive hormone-dependent regulation of autoimmunity. Science, 2013. 339(6123): p. 1084–8.

28. Guo, Y., et al., Association between Polycystic Ovary Syndrome and Gut Microbiota. PLoS One, 2016. 11(4): p. e0153196.

29. Sherman, S.B., et al., Prenatal androgen exposure causes hypertension and gut microbiota dysbiosis. Gut Microbes, 2018. 9(5): p. 400–421.

30. Le Roy, C.I., et al., Heritable components of the human fecal microbiome are associated with visceral fat. Gut Microbes, 2018. 9(1): p. 61–67.

31. Lim, M.Y., et al., The effect of heritability and host genetics on the gut microbiota and metabolic syndrome. Gut, 2017. 66(6): p. 1031–1038.

32. Goodrich, J.K., et al., Human genetics shape the gut microbiome. Cell, 2014. 159(4): p. 789–99.

33. Tremellen, K. and K. Pearce, Dysbiosis of Gut Microbiota (DOGMA)--a novel theory for the development of Polycystic Ovarian Syndrome. Med Hypotheses, 2012. 79(1): p. 104–12.

34. Klindworth, A., et al., Evaluation of general 16S ribosomal RNA gene PCR primers for classical and next-generation sequencing-based diversity studies. Nucleic Acids Res, 2013. 41(1): p. e1.

35. Fadrosh, D.W., et al., An improved dual-indexing approach for multiplexed 16S rRNA gene sequencing on the Illumina MiSeq platform. Microbiome, 2014. 2(1): p. 6.

36. Wang, Q., et al., Naive Bayesian classifier for rapid assignment of rRNA sequences into the new bacterial taxonomy. Appl Environ Microbiol, 2007. 73(16): p. 5261–7.

37. DeSantis, T.Z., et al., Greengenes, a chimera-checked 16S rRNA gene database and workbench compatible with ARB. Appl Environ Microbiol, 2006. 72(7): p. 5069–72.

38. Oksanen J K.R., Legendre P, O’Hara B, Simpson G, Henry M, Stevens H, Wagner H., Community ecology package (vegan). (2008) R Foundation for Statistical Computing, Vienna.

39. Macklin, M.T. and H.B. Mann, Fallacies inherent in the proband method of analysis of human pedigrees for inheritance of recessive traits; two methods of correction of the formula. Am J Dis Child, 1947. 74(4): p. 456–67.

40. McIver, L.J., et al., bioBakery: a meta’omic analysis environment. Bioinformatics, 2018. 34(7): p. 1235–1237.

41. Benjamini Y. H.Y., Controlling the False Discovery Rate: A Practical and Powerful Approach to Multiple Testing. Journal of the Royal Statistical Society. Series B 1995. 57(1): p. 289–300.

42. Torres, P.J., et al., Gut Microbial Diversity in Women With Polycystic Ovary Syndrome Correlates With Hyperandrogenism. J Clin Endocrinol Metab, 2018. 103(4): p. 1502–1511.

43. Zhang, B., et al., Gut Microbiota as a Potential Target for Treatment of Polycystic Ovary Syndrome. Diabetes, 2018. 67(Supplement 1): p. 2454–PUB.

44. Raygan, F., et al., The effects of probiotic supplementation on metabolic status in type 2 diabetic patients with coronary heart disease. Diabetol Metab Syndr, 2018. 10: p. 51.

45. Hall, A.B., A.C. Tolonen, and R.J. Xavier, Human genetic variation and the gut microbiome in disease. Nat Rev Genet, 2017. 18(11): p. 690–699.

46. Lerchbaum, E., et al., Adult-type hypolactasia and calcium intake in polycystic ovary syndrome. Clin Endocrinol (Oxf), 2012. 77(6): p. 834–43.

47. Aoki, R., et al., A proliferative probiotic Bifidobacterium strain in the gut ameliorates progression of metabolic disorders via microbiota modulation and acetate elevation. Sci Rep, 2017. 7: p. 43522.

48. Al Rasheed, M.R. and C.G. Senseng, Sarcina ventriculi : Review of the Literature. Arch Pathol Lab Med, 2016. 140(12): p. 1441–1445.

49. Hamer, H.M., et al., Review article: the role of butyrate on colonic function. Aliment Pharmacol Ther, 2008. 27(2): p. 104–19.

50. Lindheim, L., et al., The Salivary Microbiome in Polycystic Ovary Syndrome (PCOS) and Its Association with Disease-Related Parameters: A Pilot Study. Front Microbiol, 2016. 7: p. 1270.

